# How urban vegetation influences dynamics of *Aedes albopictus* egg density: three years of surveillance in Montpellier (France)

**DOI:** 10.64898/2026.05.15.725325

**Authors:** C. Bartholomée, C. Sutter, F. Fournet, E. Bouhsira, N. Moiroux

## Abstract

Nature-Based Solutions are increasingly promoted to address current urban challenges. While their potential effects on vector-borne disease risks have been documented, data on *Aedes albopictus*, a known arbovirus vector, remain limited in France. A previous study showed that urban vegetation moderately increases the abundance of adult mosquitoes of this species, but the monitoring period lasted only six months. Using ovitraps, we monitored *Ae. albopictus* egg density dynamics over multiple years (2022 to 2024) and analysed its environmental predictors in various urban environments. We included lagged meteorological variables, land cover metrics, and the cumulated egg densities recorded in the previous weeks as environmental predictors. Both parametric (GLMM) and non-parametric (Random Forest) models were fitted to weekly egg counts per trap. Our findings highlight that (i) egg density dynamics were related to how vegetation classes structured the landscape, (ii) growing degree days and cumulated number of eggs recorded in specific lagged time windows were the main contributors to egg density, and (iii) the non-parametric and parametric models performed similarly in terms of prediction accuracy.

## Introduction

In 2020, Nature-Based Solutions (NBS) were described by the International Union for Conservation of Nature (IUCN) as «actions to protect, manage and restore […] ecosystems, which address societal challenges […] providing human well-being and biodiversity benefits » (IUCN, 2020). That same year, NBS were promoted by the Sustainable Development Goals, notably Goal 11, which aims to « make cities and human settlements inclusive, safe, resilient, and sustainable ». Urban greening is one form of NBS, and addresses current urban challenges by mitigating urban heat islands (Emmanuel & Loconsole, 2015), increasing animal biodiversity (Threlfall *et al*., 2017), and improving human physical health (Hedblom *et al*., 2019). In 2024, Fournet *et al*. published an opinion paper highlighting the need to assess the impact of urban greening on the risk of vector-borne diseases. A scoping review conducted by Mercat *et al*. (2025) showed that urban greening and urban green infrastructures (UGIs) could increase the vector risk, depending on the vector pathogen system studied. In the case of *Aedes albopictus*, the abundance of this mosquito was found to be positively correlated with UGIs. However, only two studies on this topic had been conducted in Europe (Cianci *et al*., 2015; Manica *et al*., 2016).

*Aedes albopictus*, also known as the tiger mosquito, is an invasive species and a known vector of arboviruses, like the Dengue Virus (DENV), the Chikungunya Virus (CHIKV), and the Zika Virus (ZIKV). It has adapted to urban environments (Paupy *et al*., 2009), and uses both natural and artificial breeding sites, as well as vegetation for resting (Samson *et al*., 2013). Eggs laid during the winter months can enter a state of dormancy, or diapause, enabling them to survive desiccation and low temperatures. The species was first detected in Europe in Albania in 1979 (Adhami & Reiter, 1998) and in France in 1999 (Schaffner & Karch, 2000). By 2004, it had become established in France, and by July 2025 it had spread to 81 departments (SPF, 2025a). Since 2022, there has been an increase in indigenous arbovirosis cases, particularly in the Occitanie region (SPF, 2025a). This region is actively involved in urban greening, particularly in its two largest cities, Toulouse (Mairie-Métropole de Toulouse, 2025) and Montpellier (Ville de Montpellier, 2025). As towns in Occitanie are hotspots for arbovirus transmission and UGIs could impact the abundance of *Ae. albopictus*, it was necessary to evaluate the effect of urban vegetation on the presence and abundance of the tiger mosquito.

A first exploratory study was conducted in Montpellier between May and October 2023 by Bartholomée *et al*. (2025). They demonstrated that, although urban vegetation did not affect the probability of presence of *Ae. albopictus*, it did influence its abundance. Specifically, average patch size and cover percentage of low vegetation around traps, were found to increase its abundance. This study was the first in France to explore the relationship between urban vegetation and *Ae. albopictus*, providing initial insights into the impact of UGIs on both presence and abundance. However, the study was limited to a six-month period, and further data collection in subsequent years was needed to support more robust conclusions. Furthermore, the model used had limited predictive capacity. The objectives of the present study were, using a low-cost surveillance system: (i) exploring how urban vegetation affects *Ae. albopictus* population dynamics over an additional two years of sampling, (ii) assessing the effects of land cover and meteorological variables, and (iii) comparing the predictive performance of parametric and non-parametric models.

## Materials and methods

### Study areas

The study was conducted in Montpellier, a city located in the south of France, between 2022 and 2024 (Fig 1A). The city is composed of an impervious old city centre, urban parks, and recent residential areas with houses and private gardens on the outskirts. From 2023 to 2024, we sampled these different urban environments along a vegetation cover gradient (Fig 1B). We sampled four areas in the impervious centre (with no or minimal vegetation cover): the Diderot Institute (IMP-DID), the Bouisson Bertrand Institute (IMP-BBI), the Saint-Charles University (IMP-SCU), and the Acapulco Hotel (IMP-ACA) (Bartholomée *et al*., 2025). In 2023, we sampled four residential areas with houses and private gardens: Aiguerelles (RES-AIG), Lemasson (RES-LEM), Lodève (RES-LOD), and Soulas (RES-SOUL) (Bartholomée *et al*., 2025). In 2024, we retained only one of these residential areas (RES-LEM) and added two new ones: Hôpitaux-Facultés (RES-HOP) and Roqueturière (RES-ROQ). During both 2023 and 2024, we also sampled two urban parks: the Park of Aiguelongue (PRK-AGL) and the Botanical Garden (PRK-BOT) (Bartholomée *et al*., 2025). In 2024, we sampled the surroundings of the Institut de Recherche pour le Développement centre (IRD area, Supplementary Text 1), where our labs are based. We had access to data collected there in 2022.

**Fig 1:**
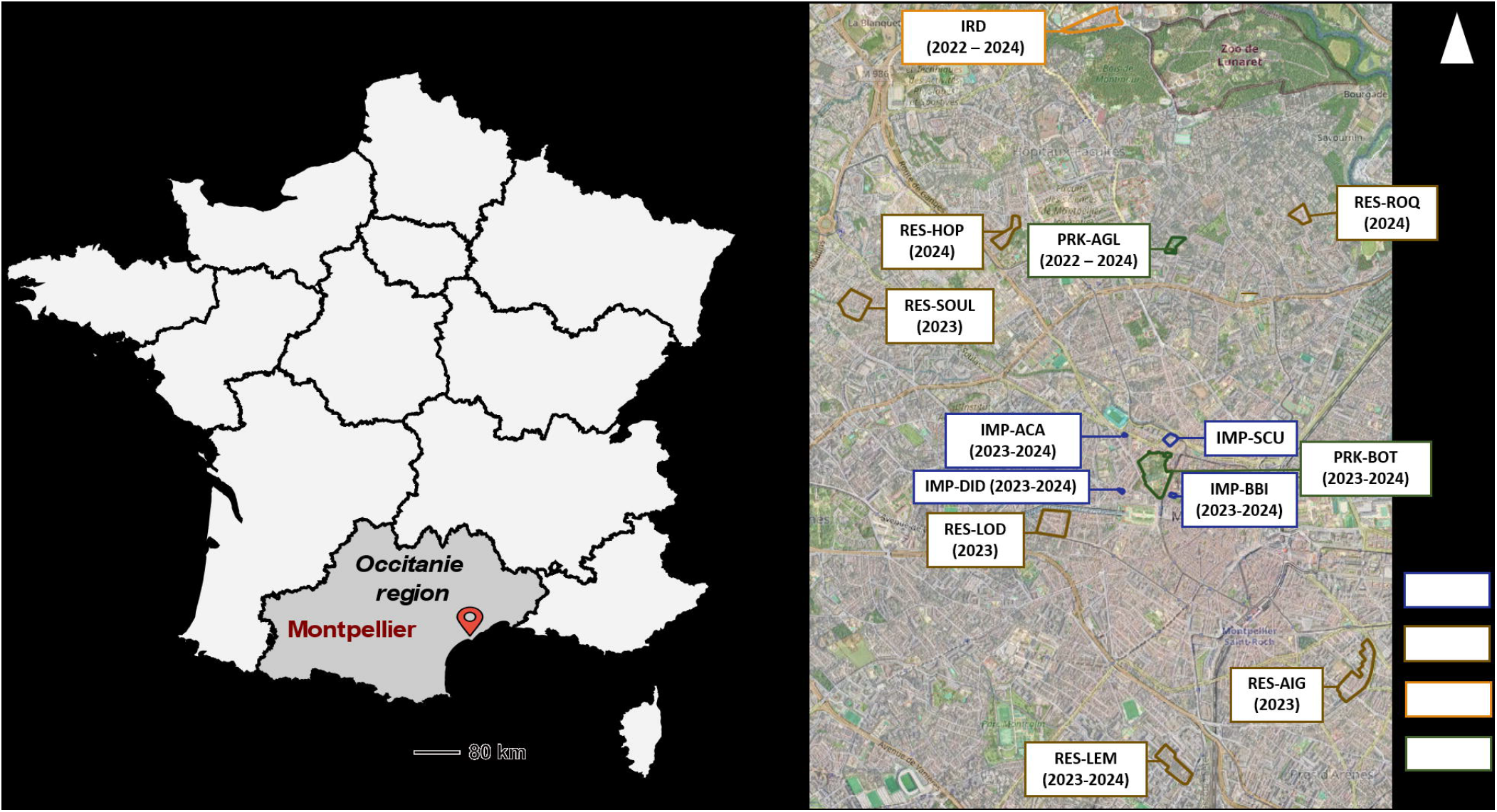
Location of Montpellier (A) (map from OpenStreetMap) and location of study areas with sampling years (B).

### Entomological sampling

We collected *Ae. albopictus* eggs using ovitraps consisting of black one-litre buckets containing 0.6 litres of clear water and containing a 10 cm × 15 cm filter paper on which the female tiger mosquitoes laid their eggs (Supplementary Figure 1). Twenty ovitraps were placed at the IRD centre in 2022 and monitored from May to September. In 2023 and in 2024, two egg traps were placed in each of the five residential areas (RES-AIG, RES-LOD, RES-LEM, RES-ROQ, and RES-SOUL) and in the Park of Aiguelongue (PRK-AGL). Due to their smaller size, only one ovitrap was placed in each impervious area (IMP-ACA, IMP-BBI, IMP-DID, and IMP-SCU) and in the additional residential area (RES-HOP). Due to their larger size, six ovitraps were installed in the Botanical Garden (PRK-BOT) and five at the IRD centre.

All traps were placed on the ground in shaded areas. Filter papers were collected, and water was replaced weekly throughout the entire study period. In 2023, twenty ovitraps were monitored from April to December: eight in residential areas, eight in urban parks, and four in impervious areas. In 2024, twenty-one traps were used over the same period: four in residential areas, eight in urban parks, four in impervious areas, and five at the IRD centre. The traps were separated by a minimal distance of 50 m, except at the IRD site in 2022, where the minimum distance was 15 m. The eggs were counted using a stereomicroscope in the IRD laboratory. *Aedes albopictus* eggs were distinguished from *Aedes geniculatus eggs*, the only other species laying eggs in ovitraps in Southern France (EID Mediterranée, 2024), using a microscope, based on the appearance of the chorion (Encinas Grandes A, 1982; Matsuo K *et al*., 1972).

### Environmental and meteorological data

To evaluate the impact of land cover on the seasonal distribution of *Ae. albopictus*, we used a detailed land cover map of Montpellier, developed in previous research (Bartholomée *et al*., 2025). This map categories land cover into five groups: low vegetation (less than 3 metres in height), high vegetation (more than 3 metres in height), roads, buildings, and ‘others’ (including railways and pathways). For each land cover category, we calculated various landscape metrics within four buffer zones around the traps (with radii of 20 m, 50 m, 100 m, and 250 m), including the percentage cover, the total edge length, and the average patch size. For the vegetation categories, we also calculated the number of patches. We used meteorological data from the station of the Observatoire Départemental de l’Eau et de l’Environnement (ODEE), located in northwestern Montpellier (Observatoire Départemental Climatologie Eau Environnement Littoral, 2024). The meteorological variables considered were related to temperature and precipitation (Da Re *et al*., 2025; Romiti *et al*., 2021). Growing Degree Days (GDD) were calculated using a daily mean temperature of 11 °C as the threshold (Neteler *et al*., 2011). All weather data were aggregated weekly: rainfall and GDD were summed, while temperature variables (mean, min. and max.) were averaged.

### Statistical analysis

The analytical framework adopted in this study is similar to those used by Bartholomée *et al*. (2025) and Taconet *et al*. (2021). The explanatory variables used in the multivariate analysis were selected based on bivariate analyses (with a p-value threshold of 0.20) and multicollinearity analyses (with a variance inflation factor (VIF) limit of 3 (Zuur *et al*., 2010)).

### Parametric models settings

This study used generalized linear mixed models (GLMMs) with a zero-inflated negative binomial distribution as parametrics models. The response variable (‘abundance’) corresponded to the number of eggs/trap/week. The models included random intercepts for sites (trap positions) nested in sampling areas. To account for temporal autocorrelation in egg counts, we incorporated an AR(1) autocorrelation structure for each sampling area and week of sampling. The AR(1) correlation structure is an autoregressive model for time-series data in which residuals are most strongly correlated at lag 1, with correlation decreasing as the lag increases (José C. Pinheiro & Douglas M. Bates, 2000).

We tested six models, with an AR(1) structure, each including the logarithm of the cumulated number of eggs laid on the same trap at different weekly time lags (from one to six weeks before sampling) as fixed effects. Model selection was based on the lowest Akaike Information Criterion (AIC). Smaller AIC values indicate models that more closely reflect the underlying reality. The best-fitting model included the logarithm of cumulated number of eggs recorded from 6 to 1 weeks before sampling as a fixed effect. AIC values for all models are reported in Supplementary Table 1.

### Bivariate analyses using GLMMs

We tested each explanatory variable individually using the GLMM structure described above. For each test, we compared the null model (without the explanatory variable) to a model including the variable, and calculated the difference in AIC (Delta_AIC_) between the two models to assess the explanatory variable’s contribution to the model’s fit. Following Burnham & Anderson (2004), a decrease of more than two AIC units was considered evidence that the variable improved the model. To facilitate comparisons between groups (land cover and meteorological variables), the AIC differences were rescaled (Delta_AICrescaled_), separately within each group, with values ranging from 0 to 1. A rescaled AIC difference close to 1 indicates that adding the variable provides the greatest improvement within its group. The time-lagged effects of meteorological variables summarised over varying time intervals were presented using cross-correlation maps (Curriero *et al*., 2005).

### Variable selection and Multivariate analyses with parametric and non-parametric models

The results of the bivariate analyses were used to select the variables for the multivariate models. Any variables that were not statistically significant in the bivariate analysis (p > 0.20) were excluded. For each meteorological or land cover variable, the time lag or spatial radius providing the greatest improvement in model fit, as indicated by the highest Delta_AICrescaled_, was retained. Finally, multicollinearity among the selected variables was assessed using VIF (Zuur *et al*., 2010).

The remaining variables were included in a parametric multivariate GLMM, and backward selection based on the AIC was applied to obtain the final, most parsimonious model. The same variables were used to create a non-parametric Random Forest (RF) (Breiman, 2001) regression model with recursive feature elimination to improve model efficiency (Speiser *et al*., 2019). Residual spatial and temporal autocorrelation was assessed for the GLMM. Spatial cross-validation was conducted for both models by training them on all areas except one and making predictions for the excluded area. To compare the models’predictions, we used the mean absolute error (MAE), calculated for all sampling areas. RF model outputs were presented as Variable Importance Plot (VIP) (Breiman, 2001) and Partial Dependence Plots (PDPs) (Friedman, 2001).

### Software and packages used

R version 4.2.3 was used (R Core Team, 2021). The ‘landscapemetrics’ (Hesselbarth *et al*., 2018) and ‘sf’ (Pebesma, 2018) packages were used to calculate the landscape metrics. GLMMs were fitted using the ‘glmmTMB’ package (Brooks *et al*., 2017) and backward selection was performed using the ‘buildMer’ package (Voeten, 2019). The ‘DHARMa’ (Florian Hartig, 2024) and ‘spdep’ (Bivand, 2022) packages were used to assess the temporal and spatial autocorrelation of the GLMM residuals, respectively. Random forest regression required the ‘caret’ (Kuhn, 2008) and ‘ranger’ (Wright & Ziegler, 2017) packages. Cross-validation was performed using the ‘CAST’ package (Meyer *et al*., 2024).

## Results

### Entomological data

A total of 71 489 *Ae. albopictus* eggs were counted by the end of 2024. As shown in Fig. 2, mosquito activity in 2023 and 2024 started mid-April and ended mid-November. There was a peak in mosquito activity in mid-June 2023 and two peaks in 2022, one occuring mid-June and one at the end of August, which were associated with peaks in rainfall.

**Fig 2:**
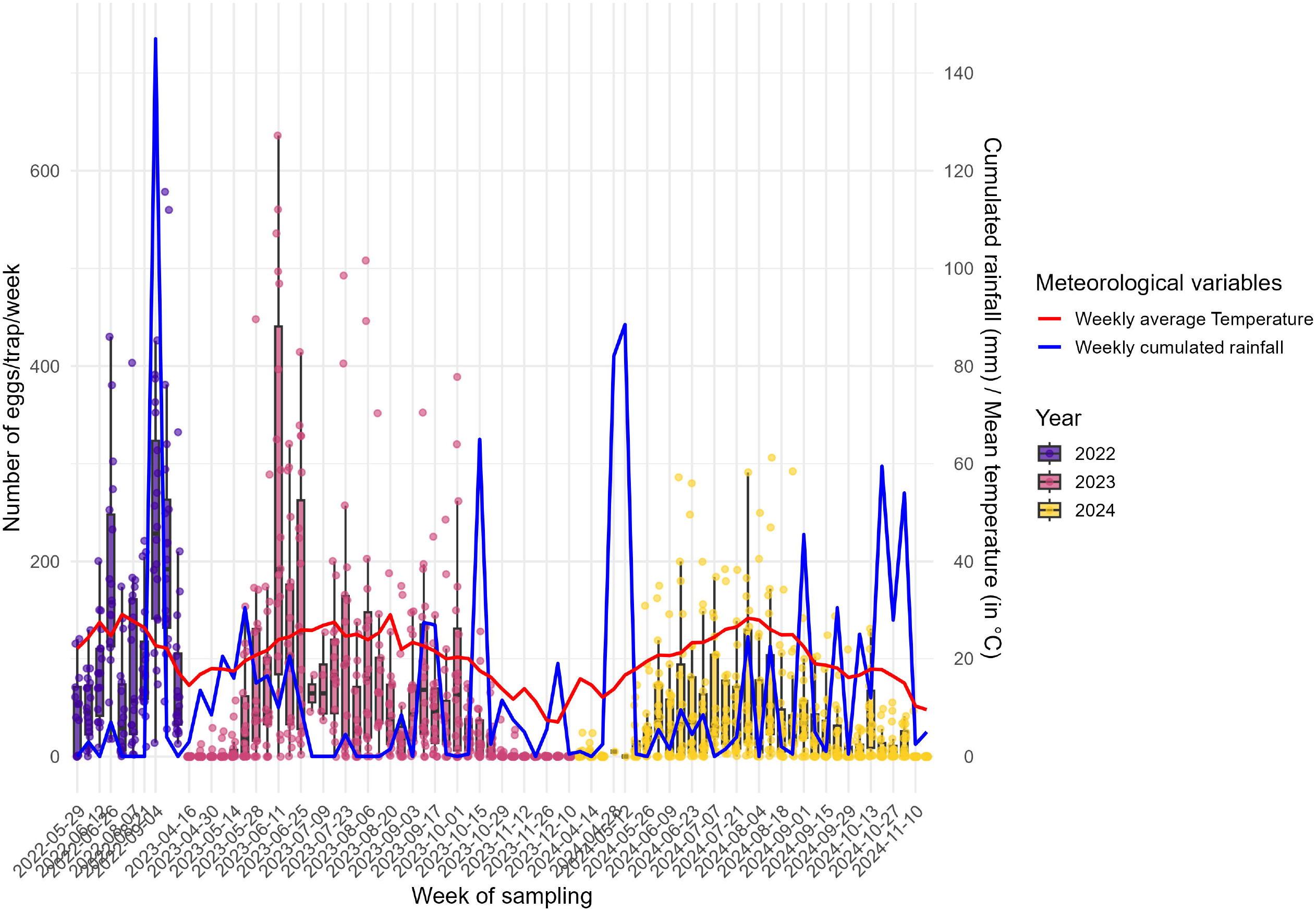
Temporal dynamics of *Ae. albopictus* eggs density per sampling week, from May 2022 to Nov. 2024, with the weekly cumulated rainfall (in mm, in blue) and the weekly average temperature (in °C, in red)

**Fig 3:**
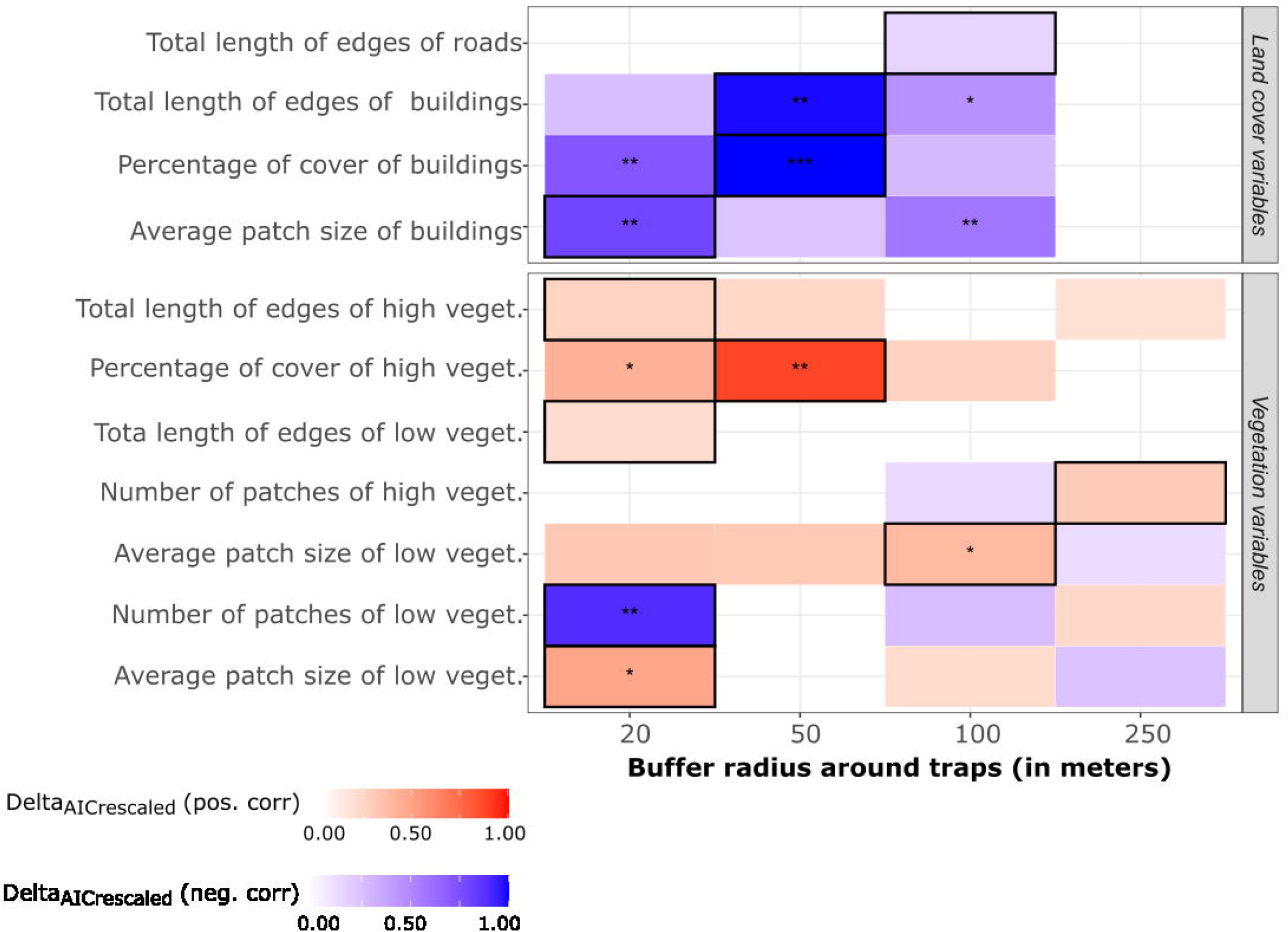
Relationships between *Ae. albopictus* eggs density and land cover variables across buffer sizes. Relationships were assessed between *Ae. albopictus* eggs number and land cover variables measured within varying buffer sizes around traps. The Delta (rescaled difference in AIC) represents the relative improvement in model fit when including a given land cover variable, compared to the null model, divided by the maximum AIC difference among land cover variable group. Values closer to one indicate stronger model improvement with that variable. Positive and negative relationships between predictors and mosquito abundance are shown separately. Boxes are colored when p < 0.2 (no asterisk: 0.05 ≤ p < 0.2; *: 0.01 ≤ p < 0.05; **: 0.001 ≤ p < 0.01; ***: p < 0.001). Box color indicates the direction of the relationship (red: positive; blue: negative), and color intensity reflects the magnitude of the difference in AIC. Abbreviations: high veget.= high vegetation, low veget.= low vegetation, pos. corr.= positive correlation, neg. corr.= negative correlation.

### Selection of explanatory variables

Fig⍰3 shows the significance and direction of the correlation of land cover variables with egg density, as well as their importance, expressed as the difference in AIC (Delta_AICrescaled_), between models including and excluding the variable. Two variables related to high vegetation were positively and significantly associated with egg abundance: the percentage of cover within a 50 m buffer around traps (Density Rate Ratio (DRR)= 1.017, 95% CI= [1.006;1.026], p-value= 0.001, Delta_AICrescaled_= 0.874), and the average patch size (DRR =1.0005, 95% CI =[1.0001;1.0010], p-value =0.037, Delta_AICrescaled_= 0.370) within a 100 m buffer around traps. Within a 20⍰m buffer, two variables related to low vegetation were significantly associated with abundance, in opposite direction: the average patch size (in m2 ; DRR= 1.001, 95% CI= [1.0002;1.0028], p-value= 0.021, Delta_AICrescaled_= 0.466) ands the number of patches (DRR =0.861, 95% CI= [0.787;0.941], p-value= 0.001, Delta_AICrescaled_= 0.894). Three building-related variables were found to be negatively and significantly associated with the number of eggs per trap per week. Within a 50⍰m buffer, these were the total edge length (in m ; IRR =0.998, 95% CI =[0.997;0.999], p-value= 0.001,, Delta_AICrescaled_= 0.998) and the percentage cover (IRR =0.975, 95% CI =[0.962;0.989], p-value =0.0005,, Delta_AICrescaled_= 1). Within a 20⍰m buffer, the average patch size (in m2) was found to be significantly and negatively associated with the number of eggs/trap/week (IRR =0.998, 95% CI =[0.997;0.999], p-value =0.002,, Delta_AICrescaled_= 0.798).

Fig 4 shows the significance (p<0.2) and importance of weeks-lagged meteorological variables in explaining egg density. When significant (red boxes), the relationship was always positive, regardless of the variable or time window considered. Rainfall and temperature data, which were summarized for the time interval three to zero weeks before sampling (box with coordinates (3,0)), best fitted egg density (highest difference in AIC relative to the corresponding null model). GDD calculated in time interval 1 to 0 weeks before sampling, best fitted egg density (Supplementary Table 2).

**Fig 4:**
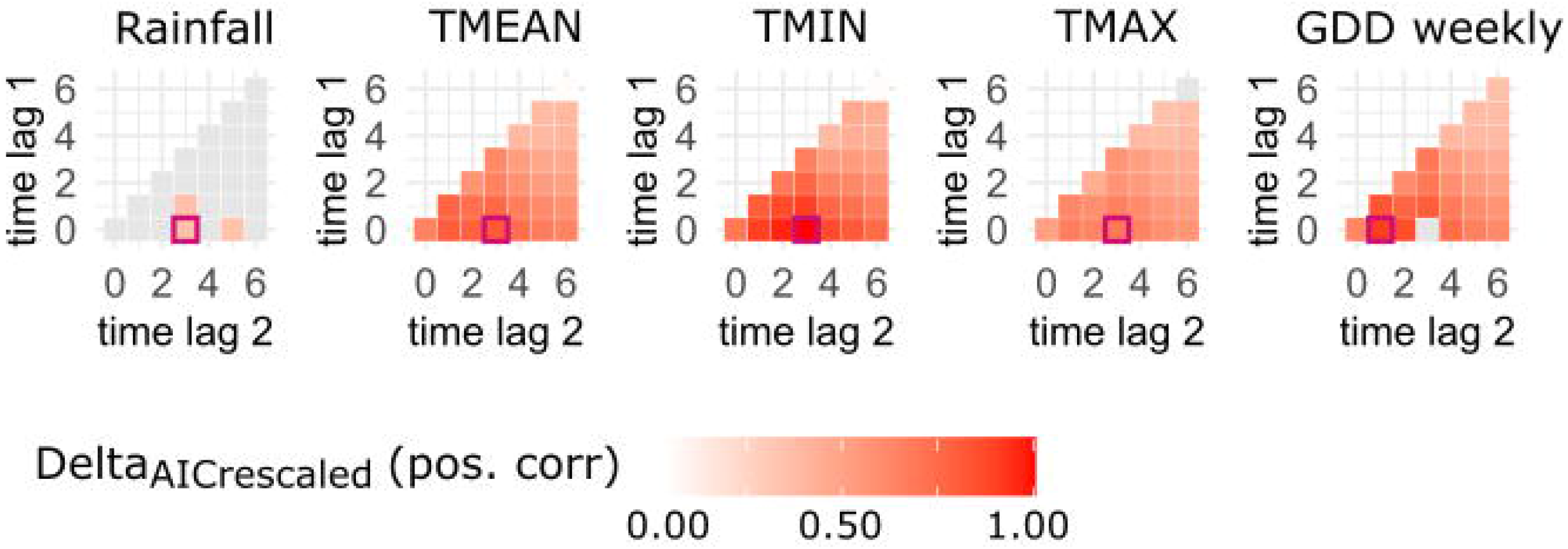
Cross-correlation maps showing an improvement in egg density models when fitted with lagged meteorological variables over varying time intervals. These lagged meteorological variables include weekly cumulated daily rainfall (Rainfall), mean daily temperature (TMEAN), minimum daily temperature (TMIN), maximum daily temperature (TMAX), and weekly Growing Degree Days (GDD). Time lags are expressed in weeks prior to sampling. The Delta_AICrescaled_ (the rescaled difference in AIC) represents the relative improvement in model fit when a meteorological variable is included, compared to the null model, divided by the maximum AIC difference among time-intervals for a meteorological variable group. Values closer to one indicate stronger model improvement with that variable. Box colour indicates the direction of the relationship (red: positive; blue: negative). Red-bordered squares highlight the time interval with the highest marginal Delta_AICrescaled_. Grey squares represent correlations with p-values > 0.2.

### Multivariate analysis: GLMM and Random Forest

Following the selection of explanatory variables, multicollinearity assessments and variable reduction steps, seven and eleven variables were retained by backward selection for GLMM and recursive feature elimination for RF, respectively (Supplementary Table 3 shows reason for variable exclusion in both models). Tables 1 and Fig 5 show the results of the parametric and non-parametric models, respectively.

**Table 1:**
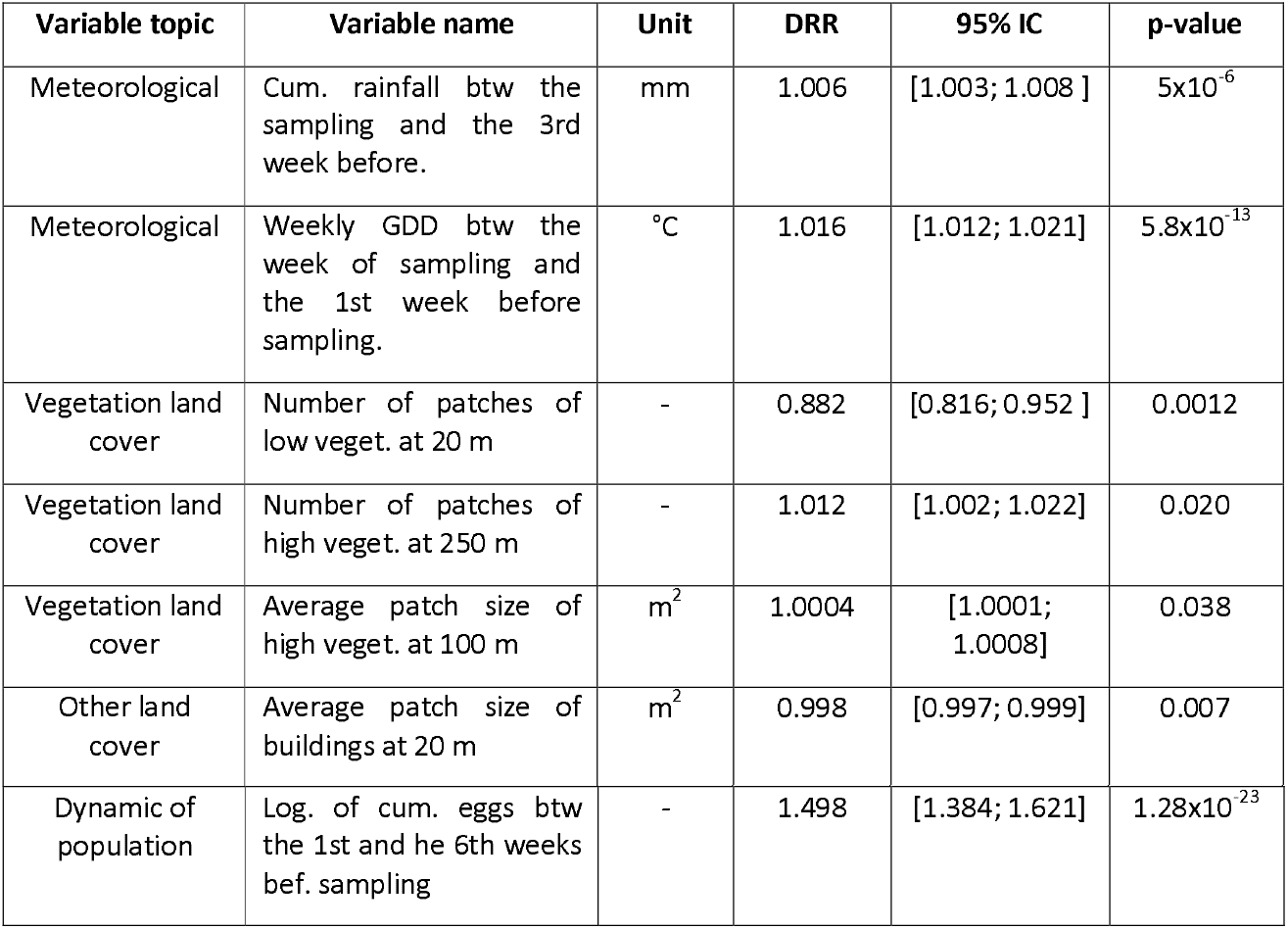
Results of the parametric multivariate analysis (GLMM) between egg density and environmental factors. The table presents the variable topic, name, unit, Density Rate Ratio (DRR), 95% IC, and associated p-value.

| Variable topic | Variable name | Unit | DRR | 95% IC | p-value |
| --- | --- | --- | --- | --- | --- |
| Meteorological | Cum. rainfall btw the sampling and the 3rd week before. | mm | 1.006 | [1.003; 1.008] | 5x10<sup>-6</sup> |
| Meteorological | Weekly GDD btw the week of sampling and the 1st week before sampling. | °C | 1.016 | [1.012; 1.021] | 5.8x10<sup>-13</sup> |
| Vegetation land cover | Number of patches of low veget. at 20 m | - | 0.882 | [0.816; 0.952] | 0.0012 |
| Vegetation land cover | Number of patches of high veget. at 250 m | - | 1.012 | [1.002; 1.022] | 0.020 |
| Vegetation land cover | Average patch size of high veget. at 100 m | m<sup>2</sup> | 1.0004 | [1.0001; 1.0008] | 0.038 |
| Other land cover | Average patch size of buildings at 20 m | m<sup>2</sup> | 0.998 | [0.997; 0.999] | 0.007 |
| Dynamic of population | Log. of cum. eggs btw the 1st and he 6th weeks bef. sampling | - | 1.498 | [1.384; 1.621] | $1.28 \times 10^{-23}$ |

**Fig 5:**
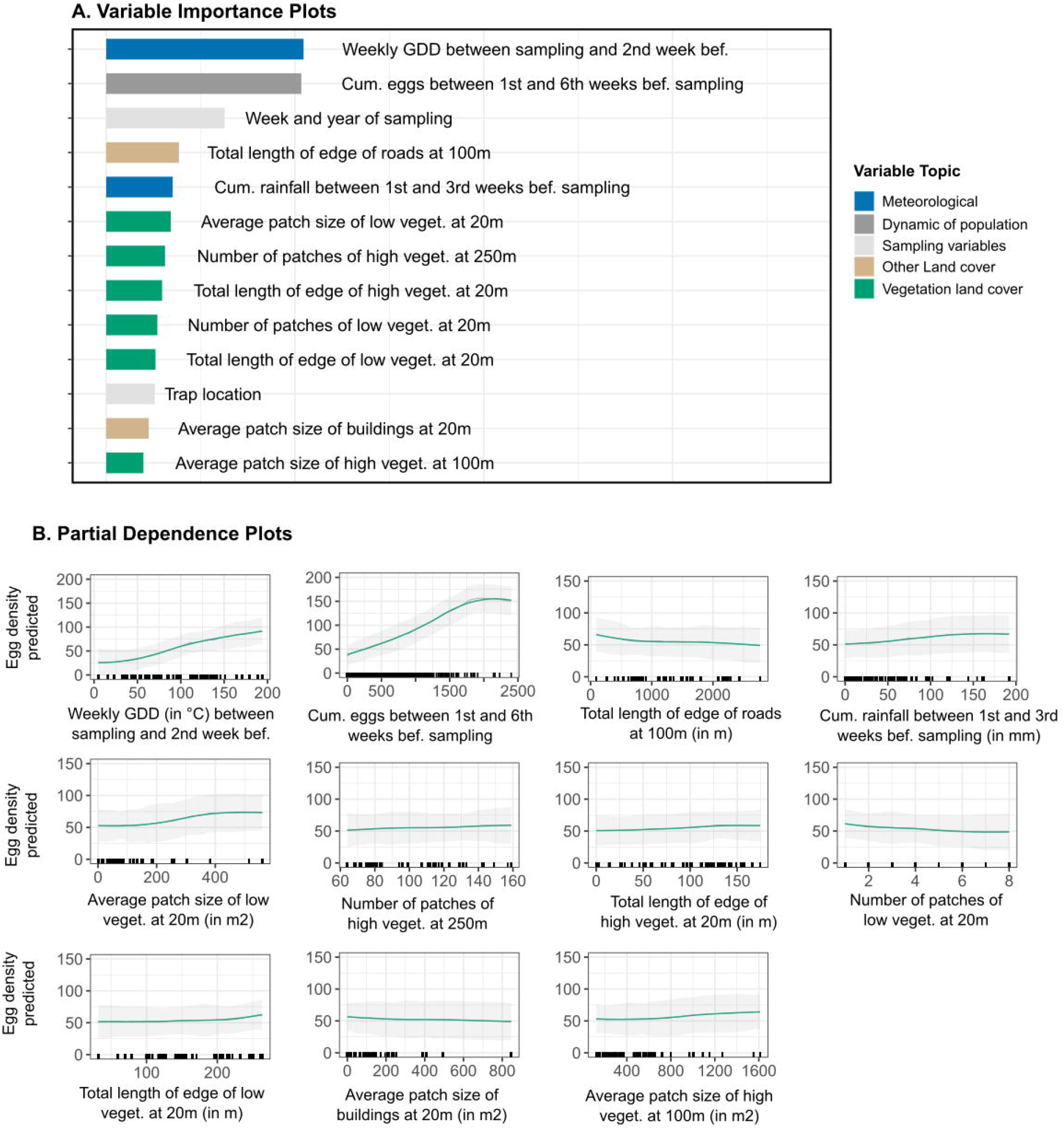
Interpretation plots for the multivariate random forest model of eggs abundance. Fig 5A is the variable importance plot (VIP). Fig 5B are smoothed partial dependence plots (PDPs) with Confidence Intervals for each selected variable in the models. The y-axis in PDPs represents the number of Ae. *albopictus* eggs/trap/week. The range of values in the x-axis represents the range of values available in the data for the considered variable. The rugs above the x-axis represent the actual values available in the data for the variable. The black line represents the unsmoothed PDP, while the green line shows the smoothed version. The shaded gray area corresponds to the 95% confidence intervals, calculated using the standard deviation and adjusted for local data density, bef.: before, veget.: vegetation, Cum.: cumulated.

According to the GLMM (Table 1), the average patch size of high vegetation within 100 m around traps was positively associated with egg density. An increase in patch size of one m^2^ corresponded to a rise of 0.04% in egg density. The number of patches of high and low vegetation showed opposite effects: the number of high vegetation patches within a 250 m buffer around the traps was positively associated with egg density (+ 1.2% per additional patch), while the number of low vegetation patches within 20 m of the traps was negatively associated with egg density (-11.8% per additional patch). The average patch size of buildings within 20m was also negatively correlated (-0.2% per additional m^2^). Meteorological variables were positively correlated with egg density: average weekly GDD in the selected time interval (1 to 0 weeks before sampling) increased egg density by 1.6% per additional 1°C, while cumulated rainfall (3 to 0 weeks before sampling) increased egg density by 0.6% per additional mm. Additionally, the logarithm of the cumulated number of eggs (6 to 1 weeks before sampling) was positively associated with egg density, with a one log unit increase corresponding to a 49.8% increase in egg density (Table 1). Compared to the null model, the GLMM improved the fit by 1161 AIC units, showed no residual deviation, or temporal or spatial autocorrelation in the residuals. Fixed effects explained 63% of the variance.

The RF model selected more explanatory land cover variables than the GLMM. However, the three vegetation-related variables of the GLMM were also included: the average patch size of high vegetation within a 100 m buffer, the number of high vegetation patches within a 250 m buffer, and the number of low vegetation patches within 50 m. Fig. 5A shows the contribution of each variable to egg density, while Fig. 5B illustrates the shape of the relationship between egg density and the explanatory variables. The first contributor was the GDD (1 to 0 weeks before sampling) with a strictly positive relationship. The second contributor was the cumulated number of eggs laid (6 to 1 week before sampling), which showed a positive association that decreased by around 2000 eggs. The third contributor was the week and year of sampling. The fourth and fifth contributors, the cumulated rainfall between 3 to 0 weeks before sampling and the total length of edges of roads within a 100 m buffer around traps, showed respectively, a slight positive and negative correlation to the density of eggs. Finally, vegetation variables moderately increased egg abundance, with the average patch size of low vegetation within a 20 m buffer having the greatest effect. The number of low vegetation patches at 20 m was slightly negatively correlated to the egg density.

Fig 6 shows the predicted number of eggs according to the multivariate GLMM and RF models, by sampling area and year. The MAE for both models is displayed. The predictions captured the temporal trends in egg abundance. However, both models underestimated abundance in IMP-ACA and overestimated it in IMP-DID during 2023 and 2024. The models showed comparable predictive accuracy, as indicated by similar MAE values. However, at the IRD site in 2022, the RF model performed better than the GLMM which overestimated densities (MAE_RF_= 70, MAE_GLMM_= 102).

**Fig 6:**
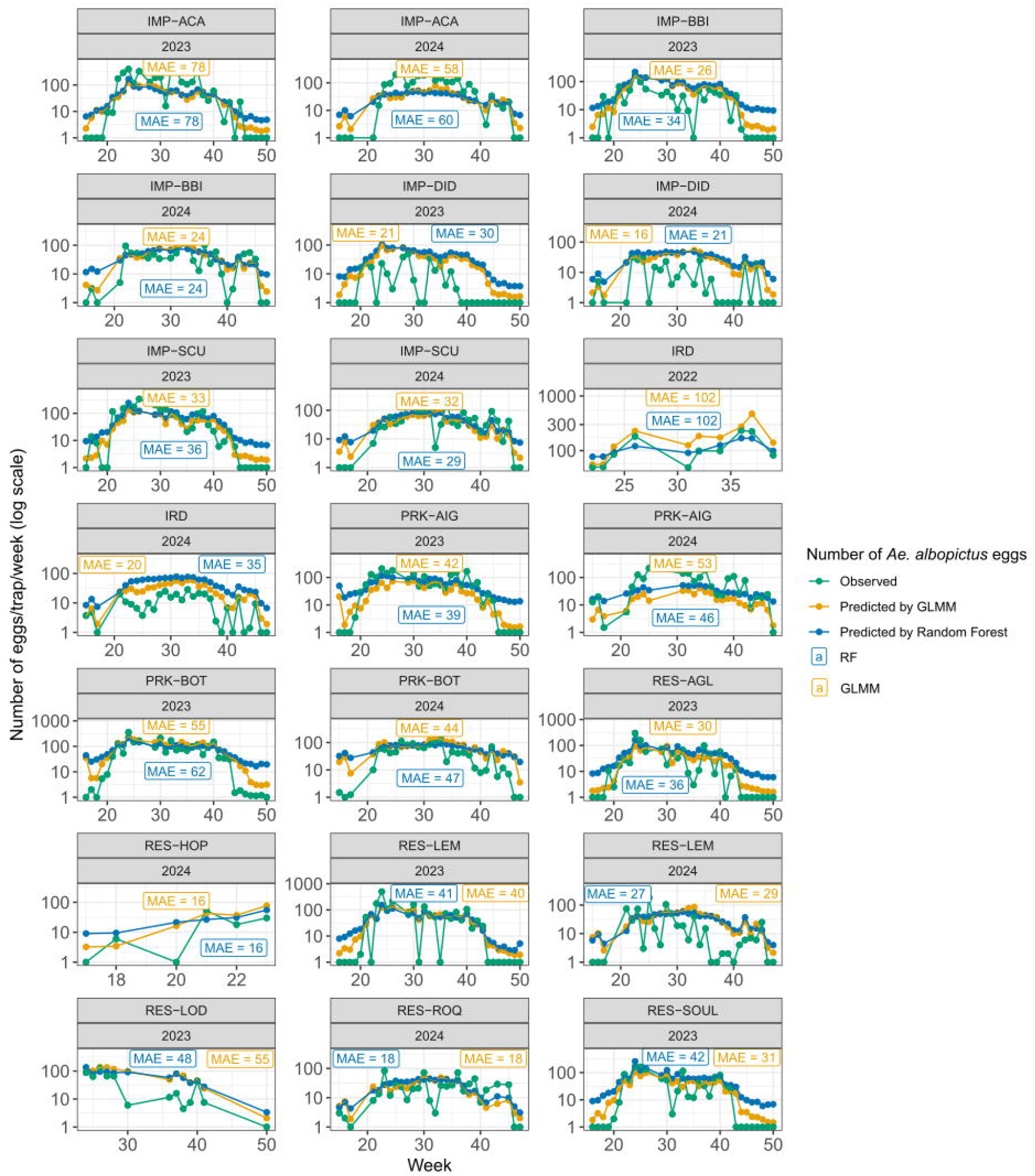
Evaluation of GLMM and RF model. The figure compares the observed abundance (in green) to the predicted abundance from the GLMM (yellow) and RF model (blue) for each out-of-sample leave-area. It also shows the Mean Absolute Error (MAE) for each model, area, and year of sampling. The y-axis represents the number of eggs per trap per week in log scale, while the x-axis corresponds to the sampling weeks. The MAE calculated for each model is displayed in yellow for GLMM and in blue for RF.

## Discussion

In this study, we analysed the dynamics of *Ae. albopictus* egg density in various urban environments in Montpellier, France, in relation to meteorological factors and the landscape, paying particular attention to vegetated areas. Both parametric and non-parametric models were used, with the past cumulated number of eggs (over the previous six weeks) acting as a predictor. This variable was found to be a significant and major predictor of the observed egg density, as previously shown by Da Re *et al*. (2025). Past egg abundance is expected to capture many slow-varying or cumulated environmental effects. Therefore, coefficients of other environmental variables (meteorology and landscape) reflect trajectories rather than raw levels and should be interpreted as immediate effects on oviposition activity, short-lag behavioural responses, or rapid ecological responses.

In Montpellier, the seasonal activity of *Ae. albopictus* began in mid-April in both 2023 and 2024, continuing until the beginning of November. This is consistent with the seasonality of *Ae. albopictus* in south-western Europe (Petrie *et al*., 2021). Regarding adult abundance in the same area (Bartholomée et al., 2025), meteorological factors were among the most important predictors of egg density dynamics in traps. The first predictor of egg density was the average weekly GDD during and just before the weeks of sampling. This reflects the known effect of temperature on life-history traits relevant to the GDD time window considered here, such as hatching rate, fecundity, and the duration of the gonotrophic cycle in *Ae. albopictus* (Delatte et al., 2009; Doeurk et al., 2024). We also found a positive correlation between egg density and the cumulated rainfall averaged over the three weeks preceding sampling. Considering this time interval, this result is consistent with the size of adult *Ae. albopictus* populations being affected by precipitation (Bartholomée et al., 2025; Roiz et al., 2015).

Both parametric and non-parametric modelling analyses showed that egg density dynamics in traps were related to landscape metrics describing the structure of the surrounding vegetated areas. Within 20 m, egg density was positively associated with the average patch size of low vegetation (under 3 m in height), but negatively associated with the number of patches in this category. This suggests that low vegetation may encourage oviposition unless the landscape is overly fragmented. A fragmented landscape may lead to a reduction in mosquito population, due to low carrying capacity of individual patches (McCormack et al., 2019). One limitation of our study is that low vegetation, as defined here (< 3m height), may comprise different types of vegetation (e.g. grass, shrubs, and hedges) that have been shown to influence egg abundance in different ways. For example, in Italy, Romiti *et al*. (2021) found that, in Lazion region, egg density was higher in areas with at least one patch of low, dense vegetation within a 100 radius, whereas Cianci *et al*. (2015) demonstrated that abundance decreased with grass cover at Sapienza University in Rome. Our work would probably have benefited from vegetation data with higher qualitative resolution. At larger distances (>= 100m), we found that egg density was positively associated with the average patch size and patch density of high vegetation (>3m height). These findings support the results of Cianci *et al*. (2015), who found a positive relationship between egg abundance and tree area within a 50m buffer zone around the traps. We also found a negative correlation between egg density and the total length of road edges within a 100 m buffer zone around the traps. This finding is consistent with the results of other studies (Romiti et al., 2021; Wang et al., 2023) and supports the hypothesis that roads act as a barrier to the dispersion of *Aedes albopictus*.

The accuracy of both GLMM and RF models in predicting egg density at sites not included in the training dataset was similar. In most cases, seasonal trends were accurately predicted. Nevertheless, there is room for improvement. Firstly, since photoperiod is one of the most influential variables affecting egg diapause (Lacour *et al*., 2015), incorporating this variable into the models would likely enhance the accuracy of predictions, particularly after September. Moreover, both models do not account for the environmental carrying capacity surrounding the traps, which may compete with them. Consequently, the model may underestimate egg densities in situations where environmental breeding sites are more attractive or abundant than traps (Barcelô et al., 2024; Fonseca et al., 2015; Huynh & Minakawa, 2022). In France, arbovirus monitoring currently begins the 1^st^ of May and ends the 30^th^ of November (SPF, 2025b). Our results indicate an earlier onset (mid-April). These findings are consistent with Caminade *et al*. (2012), who predicted that climate warming may extend the activity period of *Ae. albopictus* by approximately two weeks. This extension likely reflects both an earlier seasonal onset and a delayed cessation of activity. From a public health perspective, these results support adapting surveillance strategies by initiating monitoring earlier in the season and considering a prolonged surveillance period.

In conclusion, egg-laying dynamics in Montpellier were primarily driven by meteorological factors and varied spatially according to landscape, particularly in response to the spatial layout of vegetation. These results confirm our findings on adult abundance in the same area over several months in 2023 (Bartholomée et al., 2025). To better understand the impact of urban vegetation on the risk of *Aedes*-borne diseases, the next step would be to study other life-history traits that influence vector competence, particularly longevity.

## Acknowledgements

We would like to thank Gilbert Le Goff, Nil Rahola and Christophe Paupy for their help with the sampling design, as well as Mathilde Mercat, Coralie Grail, and Clémence Garcia-Marin for their assistance in the field and with data collection. We also thank Vincent Robert for the data he collected at IRD in 2022. Our thanks also go to the managers and staff of the various sampling sites, including the staff of the Botanical Garden. We would like to thank the Défi Clé RIVOC initiative, the Occitanie Region, and the University of Montpellier for their funding.

## Authors contribution

CB, NM and FF were involved in designing the sampling plan. CB and CS collected the field data. CB and NM performed the data analysis, and CB developed the R scripts. CB wrote the manuscript, which was then corrected by NM, EB and FF. Every co-author revised the manuscript.

