## Supplementary Files for "How urban vegetation influences dynamics of *Aedes albopictus* egg density: three years of surveillance in Montpellier (France)"

### Supplementary Information

#### Supplementary Text 1: Description of the Institut de Recherche pour le Développement (IRD)

The IRD lab is located in the North of Montpellier, adjacent to an urban forest (Bois de Montmaur), the Montpellier zoo (Zoo de Lunaret), and the green corridor along the river Le Lez. It covers an area of 3.02 ha, and it is mainly composed of buildings, parking lots, patches of shrubs and grass, and of some urban trees areas with shrubs and grass growing beneath, as shown on the following pictures. The fauna observed mainly included humans, birds (such as herons and ducks), and some domestic cats. A greenhouse and an animal facility completely enclosed, are also present for experimental purposes. There is also a permanent pool approximately 30 cm deep and measuring ten metres by ten metres.

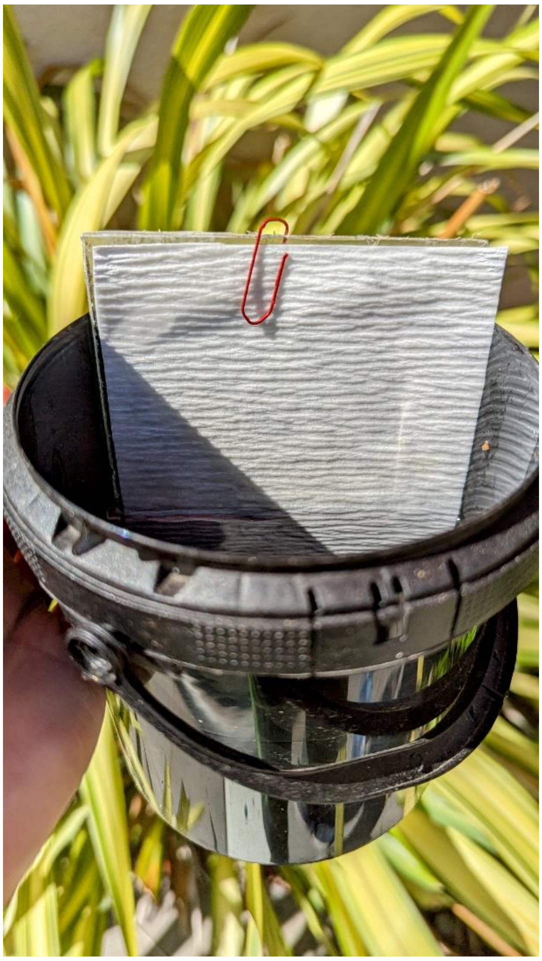
**S Fig 1: Ovitraps pictures, with the filter paper and the black 1-litre buckets (by Yseult de Dreuil)**

**S Table 1: AIC of the six models tested with the logarithm of the cumulated number of eggs at different weekly time lags (from 1 to 6 weeks before)**

| Model tested | Time lag tested | AIC |
|---|---|---|
| Model 1 | Cumulated number of eggs during the first week before sampling | 10497 |
| Model 2 | Cumulated number of eggs between the first week and the second before sampling | 10439 |
| Model 3 | Cumulated number of eggs between the first week and the third before sampling | 10397 |
| Model 4 | Cumulated number of eggs between the first week and the fourth before sampling | 10376 |
| Model 5 | Cumulated number of eggs between the first week and the fifth before sampling | 10357 |
| Model 6 | Cumulated number of eggs between the first week and the sixth before sampling | 10327 |

**S Table 2: Time-lagged meteorological variables selected based on the highest $Delta_{AICrescaled}$, with corresponding coefficients (DRR), p-values, and 95% confidence interval**

| Variable | Time-lagged selected | $Delta_{AICrescaled}$ | DRR | p-value | 95% CI |
|---|---|---|---|---|---|
| Weekly cumulated rainfall | 3 and 0 weeks before sampling | 0.334 | 1.002 | 0.122 | [0.999; 1.004] |
| Average temperature (TMEAN) | 3 and 0 weeks before sampling | 0.806 | 1.098 | $2.1x10^{-6}$ | [1.056; 1.141] |
| Minimum temperature (TMIN) | 3 and 0 weeks before sampling | 1 | 1.132 | $1.9x10^{-8}$ | [1.084; 1.182] |
| Maximum temperature (TMAX) | 3 and 0 weeks before sampling | 0.677 | 1.072 | 0.00001 | [1.039; 1.107] |
| Growing degree days (GDD) | 1 and 0 weeks before sampling | 0.903 | 1.011 | $1.18 \times 10^{-7}$ | [1.007; 1.015] |

### S Table 3: Reasons for exclusion of explanatory variables

| Variable topic | Variable code | Variable name | GLMM mutivariate analysis | | Random forest mutivariate analysis | |
|---|---|---|---|---|---|---|
| | | | Selected for the multivariate analysis (yes/no) | Reason for exclusion | Selected for the multivariate analysis (yes/no) | Reason for exclusion |
| Weeks-lagged meteorological | CUMRF_0_3 | Weekly cumulated rainfall between the sampling and the 3rd week before | Yes | NA | Yes | NA |
| Weeks-lagged meteorological | TMEAN_0_3 | Daily average temperature between the sampling and the 3rd week before | No | Correlated to TMIN_0_3, TMAX_0_3, and GDD_0_1. | No | Correlated to TMIN_0_3, TMAX_0_3, and GDD_0_1. |
| Weeks-lagged meteorological | TMIN_0_3 | Daily minimum temperature between the sampling and the 3rd week before | No | Correlated to TMEAN_0_3, TMAX_0_3, and GDD_0_1. | No | Correlated to TMEAN_0_3, TMAX_0_3, and GDD_0_1. |
| Weeks-lagged meteorological | TMAX_0_3 | Daily maximum temperature between the sampling and the 3rd week before | No | Correlated to TMIN_0_3, TMEAN_0_3, and GDD_0_1. | No | Correlated to TMIN_0_3, TMEAN_0_3, and GDD_0_1.. |
| Weeks-lagged meteorological | GDD_0_1 | Weekly cumulated Growing Degree Day between the sampling and the first week before | Yes | Although this variable is correlated with others (TMEAN_0_3, TMIN_0_3, TMAX_0_3), it is recognized as one of the key factors driving *Aedes albopictus* population dynamics (Roiz et al. 2010). | Yes | Although this variable is correlated with others (TMEAN_0_3, TMIN_0_3, TMAX_0_3), it is recognized as one of the key factors driving *Aedes albopictus* population dynamics (Roiz et al. 2010). |

| | | | | | | |
|---|---|---|---|---|---|---|
| Land cover | Patches_High_Veget_250 | Number of patches of high vegetation within a 250 m buffer zone around traps | Yes | NA | Yes | NA |
| Land cover | Patches_Low_Veget_20 | Number of patches of low vegetation within a 20 m buffer zone around traps | Yes | NA | Yes | NA |
| Land cover | %_High_Veget_50 | Percentage of cover of high vegetation within a 50 m buffer zone around traps | No | Correlated to park areas and to %_Buildings_50. VIF>3. | No | Correlated to p areas and to %_Buildings_50. VIF>3. |
| Land cover | Total_edges_High_Veget_20 | The total length of edges of high vegetation within a 20 m buffer zone around traps | Yes | Excluded during backward selection | Yes | NA |
| Land cover | Area_Patch_High_Veget_100 | The average patch size of high vegetation within a 100 m buffer zone around traps | Yes | NA | Yes | NA |
| Land cover | Area_Patch_Low_Veget_20 | The average patch size of low vegetation within a 20 m buffer zone around traps | No | Excluded during backward selection | Yes | NA |
| Land cover | Total_edges_Low_Veget_20 | The total length of edges of low vegetation within a 30 m buffer zone around traps | No | Excluded during backward selection | Yes | NA |
| Land cover | Total_edges_Roads_100 | The total length of edges of roads within a 100 m buffer zone around traps | No | Excluded during backward selection | Yes | NA |
| Land cover | %_Buildings_50 | Percentage of cover of buildings within a 50 m buffer zone around traps | No | Correlated to Total_edges_Buildings_50, %_High_Veget_50, and VIF>3. | No | Correlated to Total_edges_Buildings_50, %_High_Veget_50, and VIF>3. |
| Land cover | Total_edges_Buildings_50 | The total length of edges of buildings within a 50 m buffer zone around traps | No | Correlated to %_Buildings_50 and to Total_edges_Roads_100. VIF>3. | No | Correlated to %_Buildings_50 and to Total_edges_Roads_100. VIF>3. |
| Land cover | Area_Patch_Building_20 | The average patch size of buildings within a 20 m buffer zone around traps | Yes | NA | Yes | No |
| Land cover | Sampling environments | Sampling environments | Yes | Correlated to : for Imperviou | No | Recursive features |

| | | | | | | |
|---|---|---|---|---|---|---|
| | | | | s areas : %_High_Veget_50. For IRD: Patches_High_Veget_250 and Total_edges_Roads_100. | | elimination |
